# Differences in personality within and between five species of ants in open field tests

**DOI:** 10.1101/2023.08.16.553528

**Authors:** Alexandra Rodriguez Pedraza

## Abstract

When confronted to new situations individuals may express different kinds of behaviors. They can be atracted, explore, get immobile, attack, hide or increase their motor activity in order to confront or avoid this situation. Several studies have been conducted on vertebrate species and different patterns have been observed depending on factors as genetic or environmental ones as well as cases of rigidity or flexibility in behaviorl reaction. Less studies have been conducted on insects personalities but the current research is improving in this aspect. Here I present the case of five ant species that I tested in open field tests in order to detect if they present different response profiles when confronted to a novel environment and how these profiles can vary depending on factors as species, ambient conditions or ecological realities of the studied populations. In this article I expose hypothesis to explain you how they react in these circumstancies and how the observed differences can subtend some of their living realities.

## Introduction

Personality, neophobia and temperament are three concepts that have been more and more used by psychologists and ethologists (Bates 1986, Chapple et al. 2012) to describe the interindividual differences in the expression of the behavior present early in individuals’ lives and stable across the time (Bates 1989, Goldsmith et al. 1987). Personality can include aspects of behavioural responses as reactions towards predators, neophobia and social behaviour towards conspecifics. These behavioral characteristics may condition dispersal behavior, approach towards novel objects and novel food items consumption (Sinn et al. 2008).

The literature concerning interindividual differences in personality is very extensive and some studies converge to say that constancy exists in species like horse and salamander (Le Scolan et al. 1997, Sih et al. 2003) whereas other studies found that there is no constancy across the contexts in species like pumpkinseed sunfish and prairie voles (Coleman and Wilson 1998, Neff and Sherman 2004, Lee and Tang-Martinez 2009). These different observations vary depending on the studied species, on the methods that are used and on the behavioural aspects that are explored.

Few studies have been conducted on reptiles, however, differences in temperament have been observed in lizards for example and these differences appeared as being involved in learning processes (Carazo et al. 2014). Actually, in this case, bold individuals appeared to learn faster than shy ones because of their tendency to explore earlier the objects and the experimental apparatus. In the Namibian rock agama, *Agama planiceps*, it has been demonstrated that boldness can induce dangerous situations as bold individuals are more easily trapped than shy ones (Carter et al. 2012)

Neophobia is one of the aspects of personality and it consists in the fear towards novel stimuli or novel situations. First, a stimulus can be novel because it has simply never been experienced by an individual during its lifetime (Brown et Chivers 2005). This phenotype has recently received much ecological attention, primarily in the context of decision making and even in large scale behaviors as migration. For example, low levels of neophobia were reported in migratory Sardinian warblers, *Sylvia melanocephala momus*, when compared to garden warblers, *Sylvia borin* (a non-migratory species) (Mettke-Hofmann et al. 2005). Furthermore, in a study conducted by Candler and Bernal (2015), differences of boldness were observed in cane toads as individuals from native populations did not approach a novel object while more than half of the individuals from introduced populations did. It has also been reported that early experience with novel objects in laboratory environments can result in low neophobia levels in young hand reared parrots *Amazona amazonica* compared with individuals raised by their parents in the simpler nest box environments that present lower objects diversity (Fox and Millam 2004).

Personality and temperament have been studied in several vertebrate species and it has been observed that different factors influence their expression, from genes to environmental conditions (Tremmel and Muller 2013). Personality has also been studied in invertebrates as in the crayfish *Pacifactacus leniusculus* (Galib et al. 2022) but it has been little studied in insects.

Ants are one of the most represented taxa in the world. This group contains a huge number of species with a broad diversity of phenotypes and behavioural patterns. This enormous biological diversity has led to the existence of ants in several ecosystems and biotopes.

The success of ants in the world is mainly due to their social organisation in castes and their anatomical specialisation thanks to the existence of morphotypes as queens, workers and soldiers that present entirely different phenotypes (Chitkka et al. 2012). Most of the literature in ants explores the physical differences in phenotypes and anatomy to characterize individuals and to explain many of the biological phenomena observed in this taxon. However, no studies have been conducted on personality differences between castes.

In this work I explored some aspects of the personality of ants in order to understand what factors can explain their behavioral diversity and plasticity as well as their tendency to be good candidates for biological invasions.

## Material and methods

### Experimental protocol

Open field tests have been used from decades to study exploration in vertebrates like rats. These tests measure exploratory locomotor behavior and general activity in rodents.

Since its introduction, 40 years ago, the open-field test has become one of the most widely used instruments in animal psychology to test aspects of personality. The open field test is a common measure of exploratory behavior and general activity. The open field is an enclosure generally square, rectangular or circular with surrounding walls that prevent escape (Gould et al. 2009).

Originally introduced as a measure of emotional behaviour in rats, open field exploration has proven to be equally successful with mice. The test offers the possibility to assess novel environment exploration, general locomotor activity, and some aspects of behaviour in rodents. When placed in an open field, animals may display different behaviours. Some wander through the area to explore the new environment. Rodents will typically spend a significantly greater amount of time exploring the periphery of the arena, usually in contact with the walls than the unprotected centre area (Taiwo 2015). Open spaces induce fear behaviour in animals that have an aversion to unfamiliar environments, and these animals will seek dark regions or protective crevices. Rodents that spend significantly more time exploring the unprotected centre area are considered bolder and less anxious than individuals that explore in close contact with the walls.

In this study I adapted the classical open field protocols from rodents studies to the study of ants personality.

For this aim I compared five species of ants in open field exploration tests. For this test I used plastic white opaque glasses of 7 oz.

Four ant species (*Atta cephalotes, Crematogaster crinosa, Solenopsis invicta, Camponotus atlantis)* were tested from four different sites in Colombia (South America) and one species (*Messor barbarus*) was from a European site in France.

#### Location of the ant species

*Atta cephalotes*: Cali Colombia

*Crematogaster crinosa*: Punta Roca (Barranquilla) Colombia

*Solenopsis invicta*: Islabela Island (Colombia)

*Camponotus atlantis*: Cartagena Colombia

*Messor barbarus*: Luminy-Marseille (France)

### Biological material

I present below the different species tested for this study.

Messor barbarus are hard worker harvester ants found in Southern Europe and Northern Africa and they are the most common ants in arable fields in northeastern Spain (Westerman et al. 2012).

This species is found to act in accordance with the optimal foraging theory, which predicts that selectivity in ants increases with increasing richness of resources in an area, as well as with increasing distance from starting location. Highly frequented trails have a higher mean rate per worker, meaning the harvesters returned higher rates of resources more efficiently along these trails. These trails draw more foraging ants to retrieve seeds on the whole, and the foraging ants return seeds at a higher rate per capita (Detrain et al. 2000). This foraging pattern indicates that relative food abundance along paths may impact the patterns of trail foraging behavior in *Messor barbarus*. For trails spanning long distances, ants exhibit behavior of strong chemical marking on preferred seeds to allow for the creation and maintenance of the route. However, in this study I wondered if ants’ personality could also have an influence on choosen trails.

For testing this hypothesis I first monitored a *Messor barbarus* colony and noted the number of ants forraging at different distances from the nest in order to identify potential patterns of foraging. Then, I tested individuals taken from two different distances from the entrance of the nest and conducted the open field test on these different individuals in order to determine if they presented different behavioral patterns.

*Atta cephalotes* is a leafcutter ant from the tribe Attini widely distributed from Mexico to Brasil (Correa et. Al 2005). In Cali, Colombia, it is considered as an invasive species as it has widely colonized almost all the city and big colonies can be observed that have even destroyed some soil support in several neighborhoods (Montoya-Lerma et al. 2006).

Crematogaster crinosa is a Neotropical ant species found across a range of habitats from southern Texas to Argentina but the largest colonies are found in dryer areas such as seasonally dry forest. They are difficult to distinguish and workers may not always be clearly identified. In Costa Rica and Colombia mangrove forests are dominated by Crematogaster azteca and Crematogaster crinosa (Longino 2003). The individuals tested here were observed at Punta Roca locality close from Barranquilla in a dry rocky área near to the beach (at 100m from the coast).

*Solenopsis invicta* is a native ant from tropical and subtropical South America that has obtained international notoriety as it became an enormously successful invasive ant throughout much of the southern United States, being one of the 100 worst invasive species in the world (IUCN/SSC Invasive Species Specialist Group). It is now spreading rapidly in parts of the Caribbean, and new infestations have been detected and exterminated in Arizona, California, Australia, New Zealand, and southern China. The probability of new invasions is therefore quite high and *S. invicta* must be considered a potential threat worldwide in all areas where climates are suitable (Helmes et al. 2017).

I tested 30 individuals of *Solenopsis invicta* living at Islabela, an island which is situated in the Caribean see in front of Cartagena City. I compared the open field test reactions of individuals captured close to the nest entrance and individuals captured far from the nest entrance.

*Camponotus atlantis* are commonly found under rocks next to and under Acacia trees (Sharaf et al. 2017). This species of ants is widely spread in the world. *Camponotus* is an extremely large and complex, globally distributed genus. At present, more than 1000 species and nearly 500 subspecies belonging to 45 subgenera are described (Bolton, 2012) and it could well be the largest ant genus of all. The enormous species richness, high levels of intraspecific and geographic variation and polymorphism render the taxonomy of *Camponotus* one of the most complex and difficult. These ants live in a variety of habitats and microhabitats and the large size of the genus makes any characterisation of their biology challenging. Nests are built in the ground, in rotten branches or twigs, or rarely into living wood (Bolton, 1973) and most species possess a highly generalistic diet (García et al. 2013).

For this study I tested 32 individuals of *Camponotus atlantis* from Cartagena (Colombia) in the hot external place (at 34°C) of the gardens of Santa Clara Hotel and 23 individuals from the same Hotel in Cartagena in the warm conditions inside the hotel (24°C).

### Experimental procedure

For each test I put the ant into a clean plastic glass and I recorded its behaviour during 3 minutes. The following behavioural acts were recorded and counted.

**Table 1.** Behavioural items observed during the open field tests.

| **Behaviour** | **Description** |
| --- | --- |
| **Walks** | The ant walks on the bottom of the glass |
| **Climbs** | The ant climbs on the walls of the glass |
| **Run around the open field** | The ant runs around the central part of the glass without crossing it |
| **Antenna grooming** | The ant cleans its antenna |
| **Cross the open field** | The ant runs trough the central part of the glass leaving the walls of the glass |
| **Explores the area** | The ant walks and touches the plastic glass with its antennas |
| **Total marching acts** | Walks+Climbs |
| **Immobility** | Time spent by the ant in an immobile position |

### Experimental conditions

- ***Comparison of the number of individuals foraging close and far from the nest entrance*** I conducted field observation and counting on the activity of *Messor barbarus* ants at Luminy University Campus in southern France close to Marseille. I counted the number of ants that foraged at different distances from the entrance of the nest between 6 and 7pm as it is the moment of higher activity in these ants that avoid hot sunny conditions during the day and stay into the nest. The results of this counting are presented on **Figure 7**.
- ***Comparison of neophobia between ants taken close from the nest entrance and ants taken far from the nest entrance*** I conducted this experiment with the *Solenopsis invicta* and *Camponotus atlantis*. At Islabela Island I tested *Solenopsis invicta* ants taken from the entrance of the nests and taken more than 2 metters from the nest entrance. At Cartagena city, I used the same protocol to study *Camponotus atlantis* but in this case I also tested for the temperature factor on the individuals’ reaction. So, for this experiment I tested a group of *Camponotus atlantis* outside the hotel at 34°C and into the hotel where air conditioning was on (24°C).
- ***Comparison of neophobia between soldiers and workers*** For the big species (*Atta cephalotes* and *Messor barbarus*) that present clearly define castes of soldiers and workers I compared the reaction of individuals from these different castes at a same temperature.
- ***Ants’ manipulation*** For each ant tested in this study the ant was put on the glass, tested and then put in another plastic container while other ants were tested in order to avoid testing two times the same ants. At the end of the experiments the ants were released at their original place of capture.

**Figure 1.**
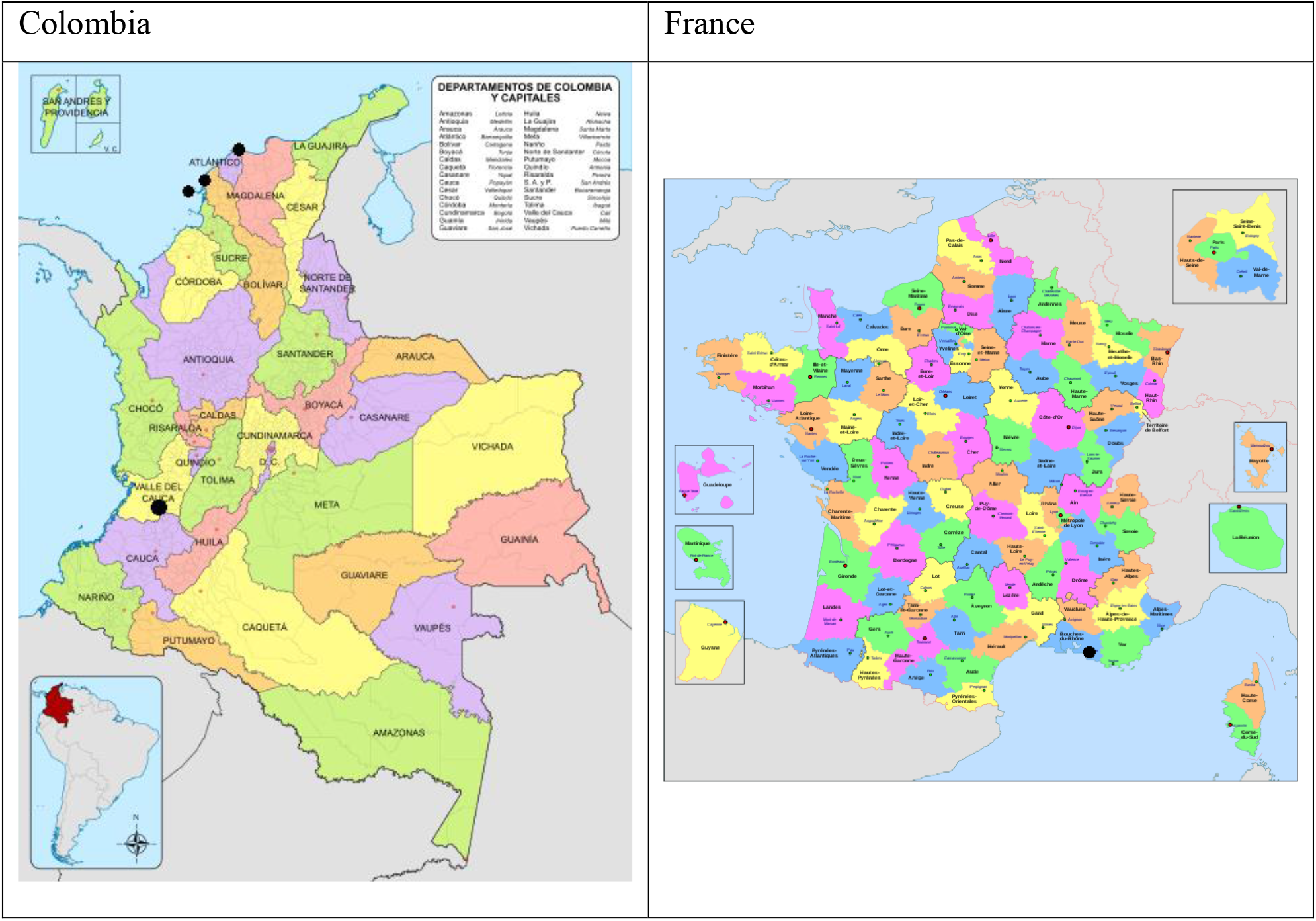
Location of the ant colonies that were selected for the study.

**Figure 2.**
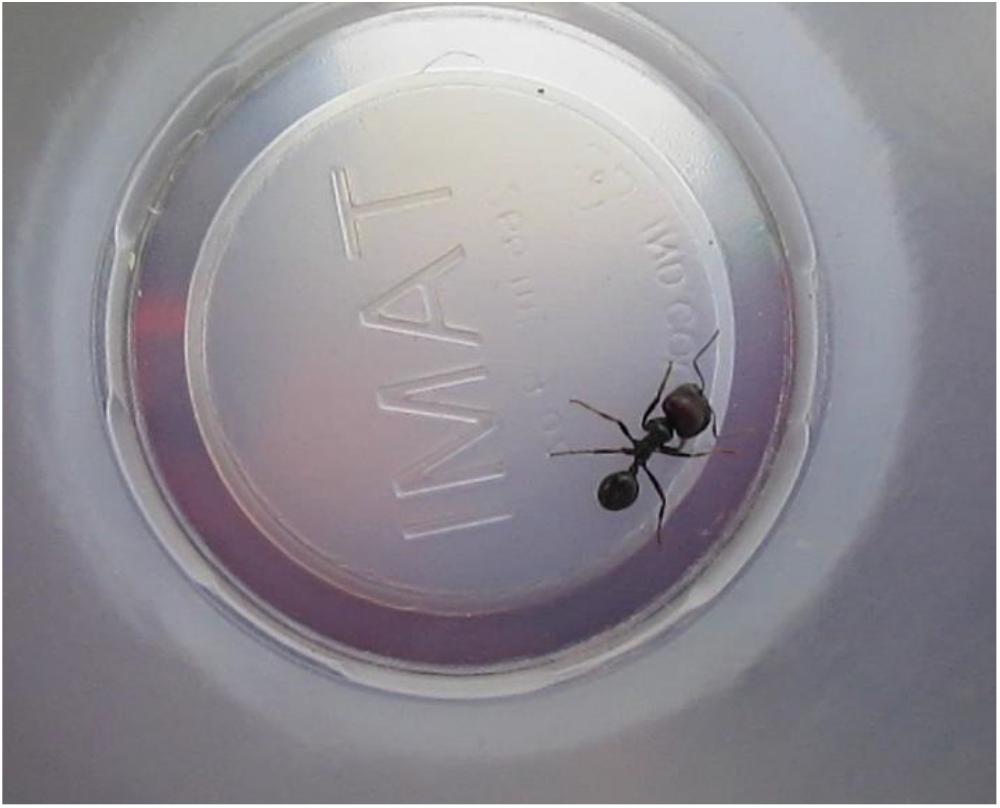
Messor barbarus.

**Figure 3.**
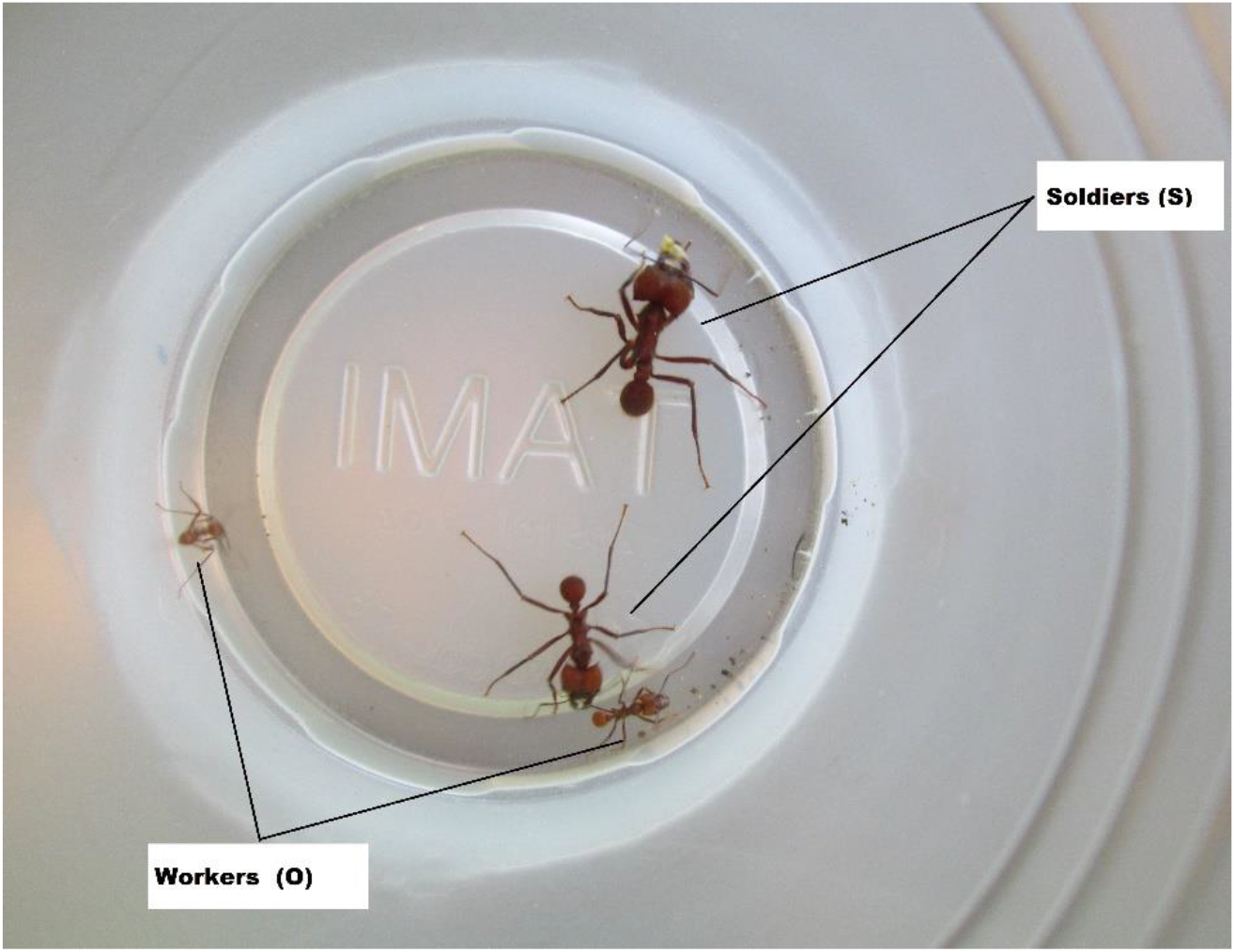
Atta cephalotes. Four individuals are presented here for the comparison of castes’ sizes

**Figure 4.**
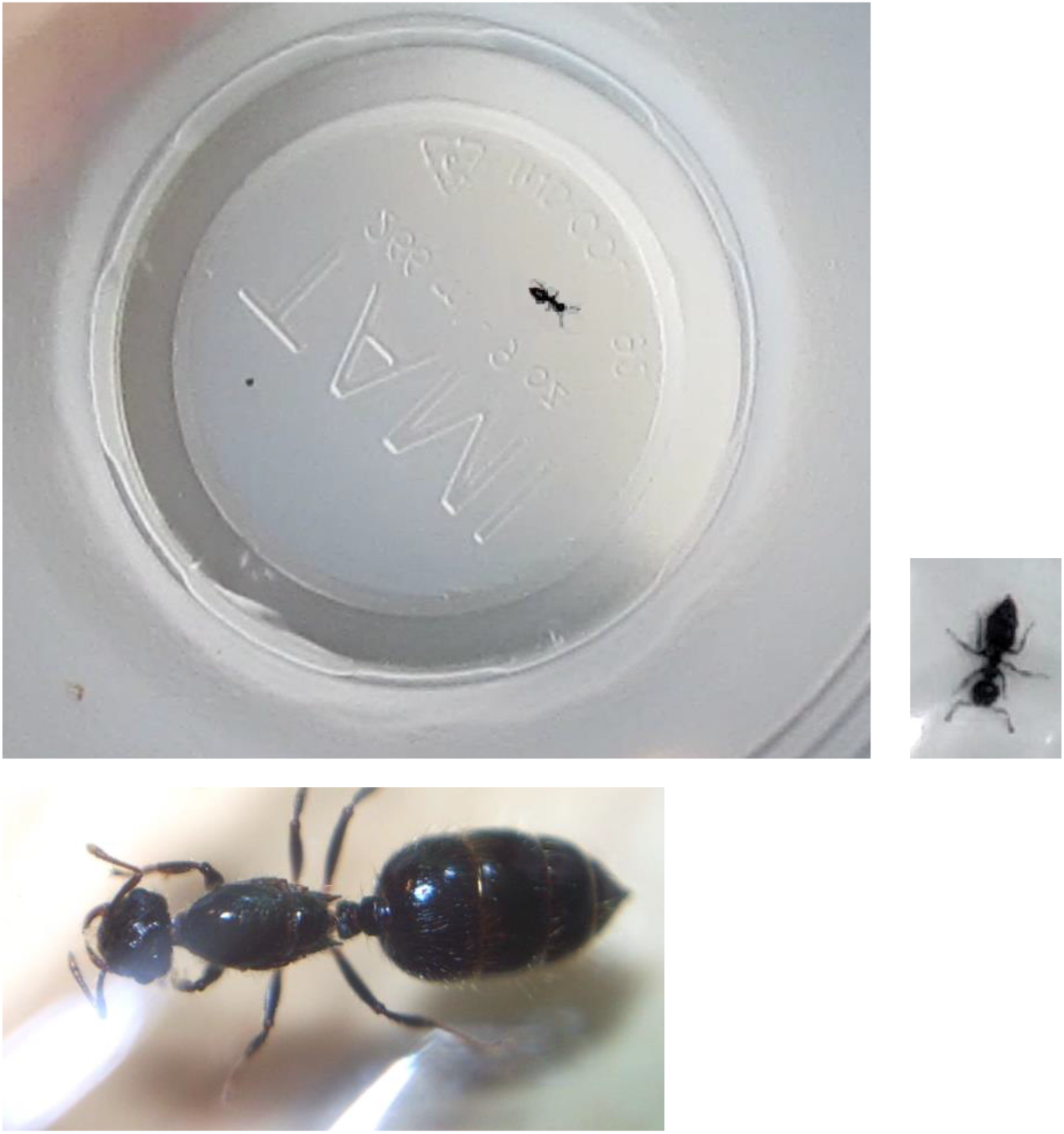
Crematogaster crinosa.

**Figure 5.**
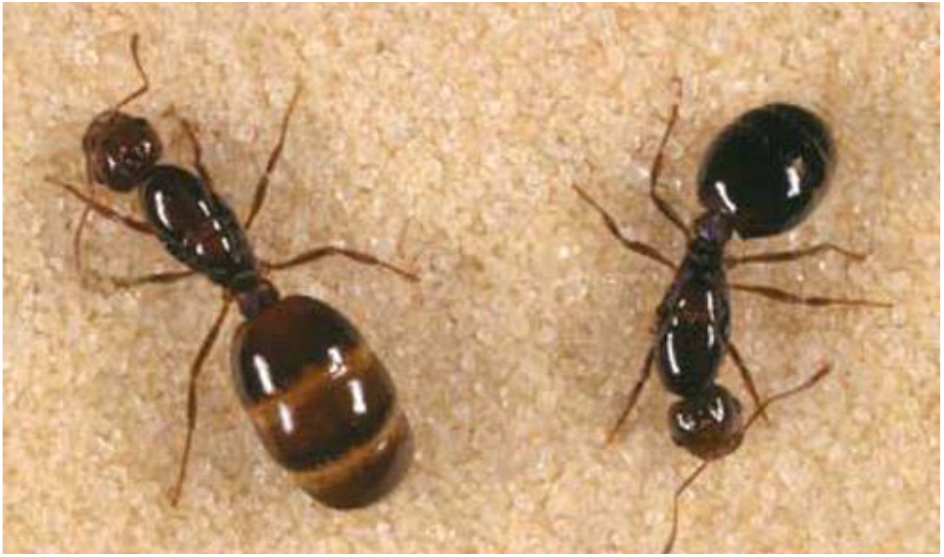
Solenopsis invicta.

**Figure 6.**
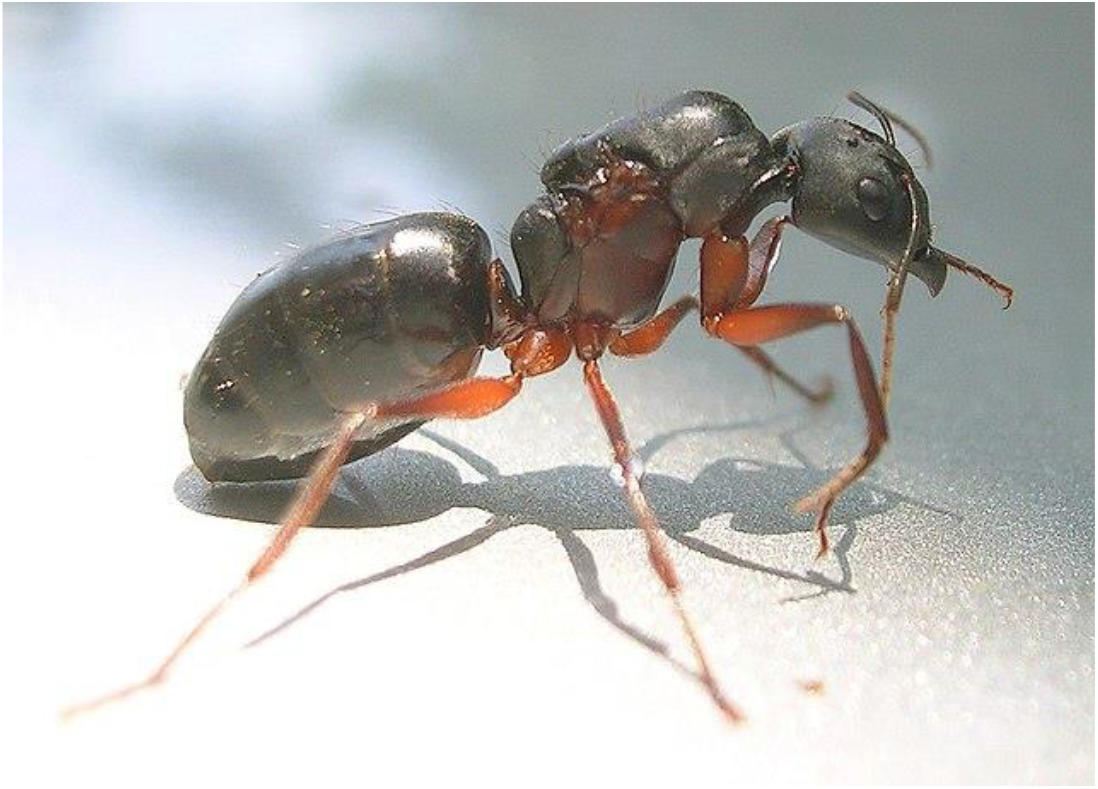
Camponotus atlantis.

**Figure 7.**
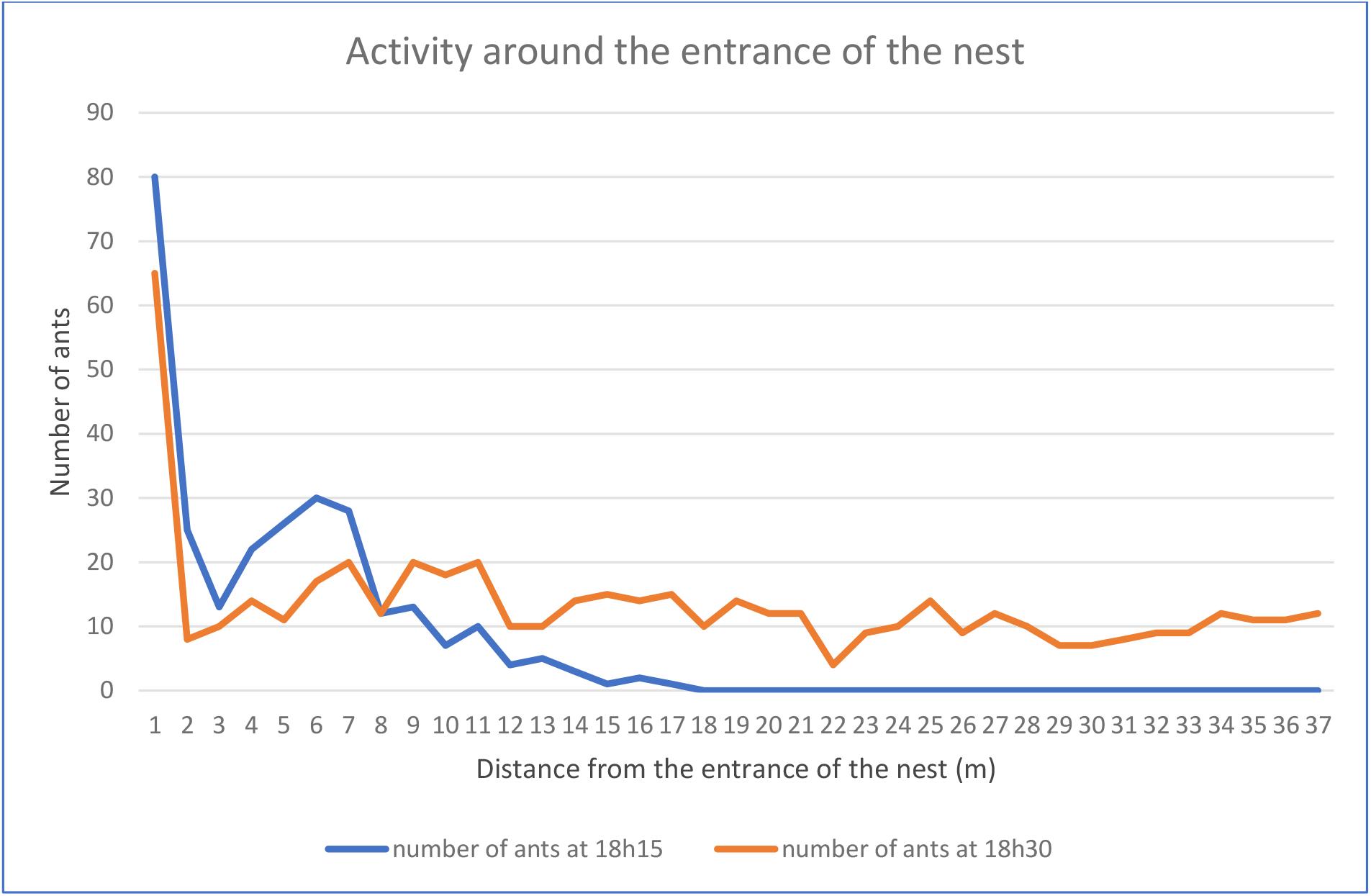
Foraging activity of *Messor barbarus* at different distances from the nest.

### Statistical Analysis

As sampling sizes were little, I conducted Mann and Withney non parametrical tests to compare ants’ different behavioural responses between:

- Hot and warm conditions
- Workers and Soldiers
- Individuals taken close from the nest entrance (low explorers) and far from the entrance (big explorers).

My aim was to know if temperature could have an influence in neophobia reaction towards a novel environment and to know if the personalities were different depending on the ants’ status (workers vs soldiers and big explorers vs low explorers).

## Results

In the observations conducted on *Messor barbarus’* activity around the nest during the big foraging activity period of the day (6pm-7pm) I noted that there was a majority of ants foraging in a circle of two meters around the nest entrance (almost 60% of the foraging individuals) **(Figure 7)**. The other 40% of individuals was distributed further (around 20% between 2 and 12 meters far from the entrance of the nets and 20% between 12 and 37 meters from the nest).

In the tests conducted on *Solenopsis invicta* at Islabela Island I observed that individuals captured far from the nest entrance climbed significantly more frequently the glass walls in the open field test than the individuals captured close to the nest entrance. They also crossed the open area of the open field more frequently (p<0.01) than the individuals captured close to the nest entrance **(Figure 8)**.

**Figure 8:**
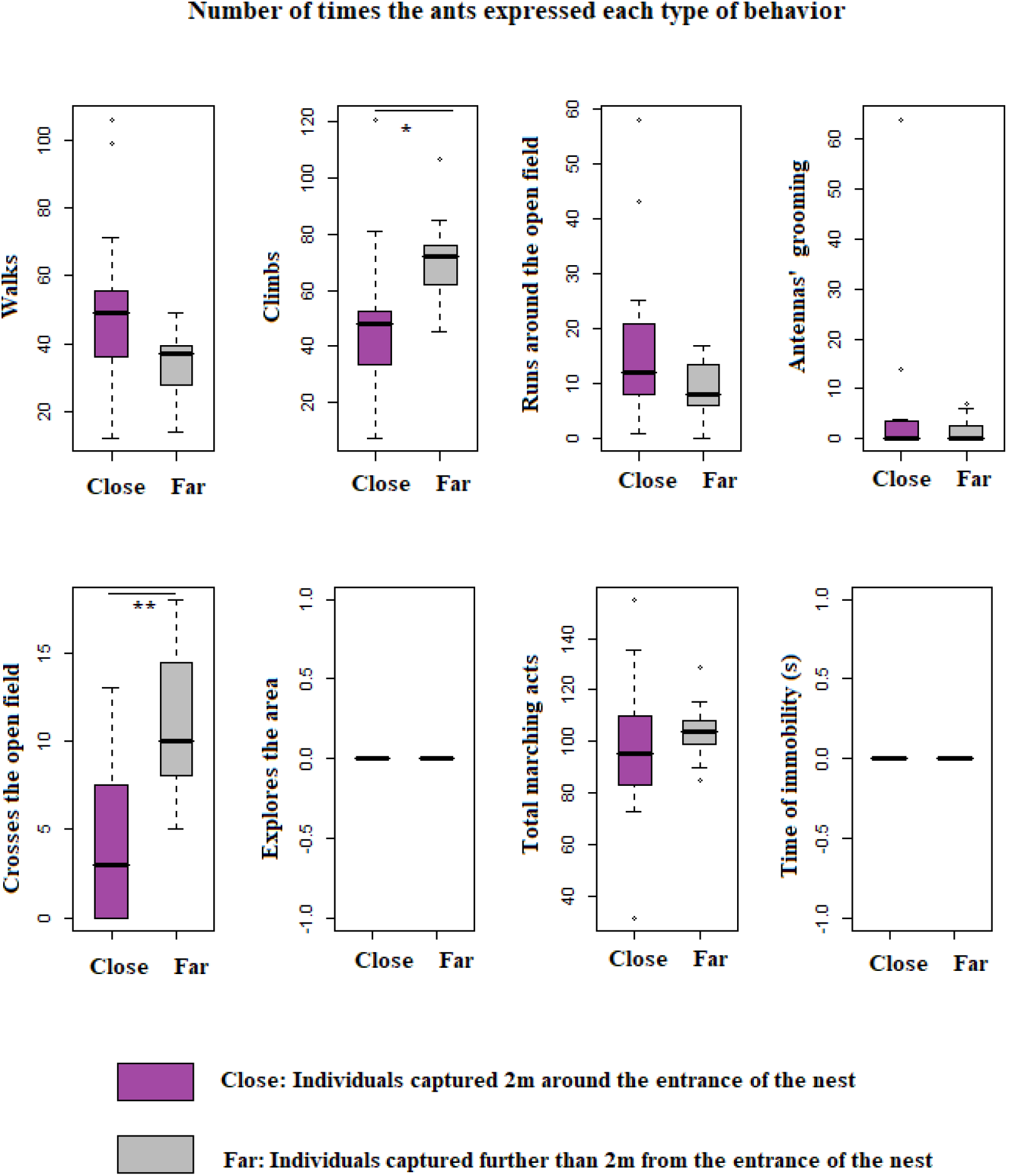
Comparison of 30 *Solenopsis invicta* activity when captured close and far from the nest in Islabela Island.

In the test conducted on *Camponotus atlantis* comparing hot versus warm (cold) conditions, I observed that individuals put in hot conditions at 34°C walked, explored and run around the open area significantly more frequently than individuals tested at 24°C **(Figure 9)**. However, they explored the glass with their antennas more frequently at 24°C than at 34°C. They spent significantly more time in immobile positions when tested at 24°C.

**Figure 9:**
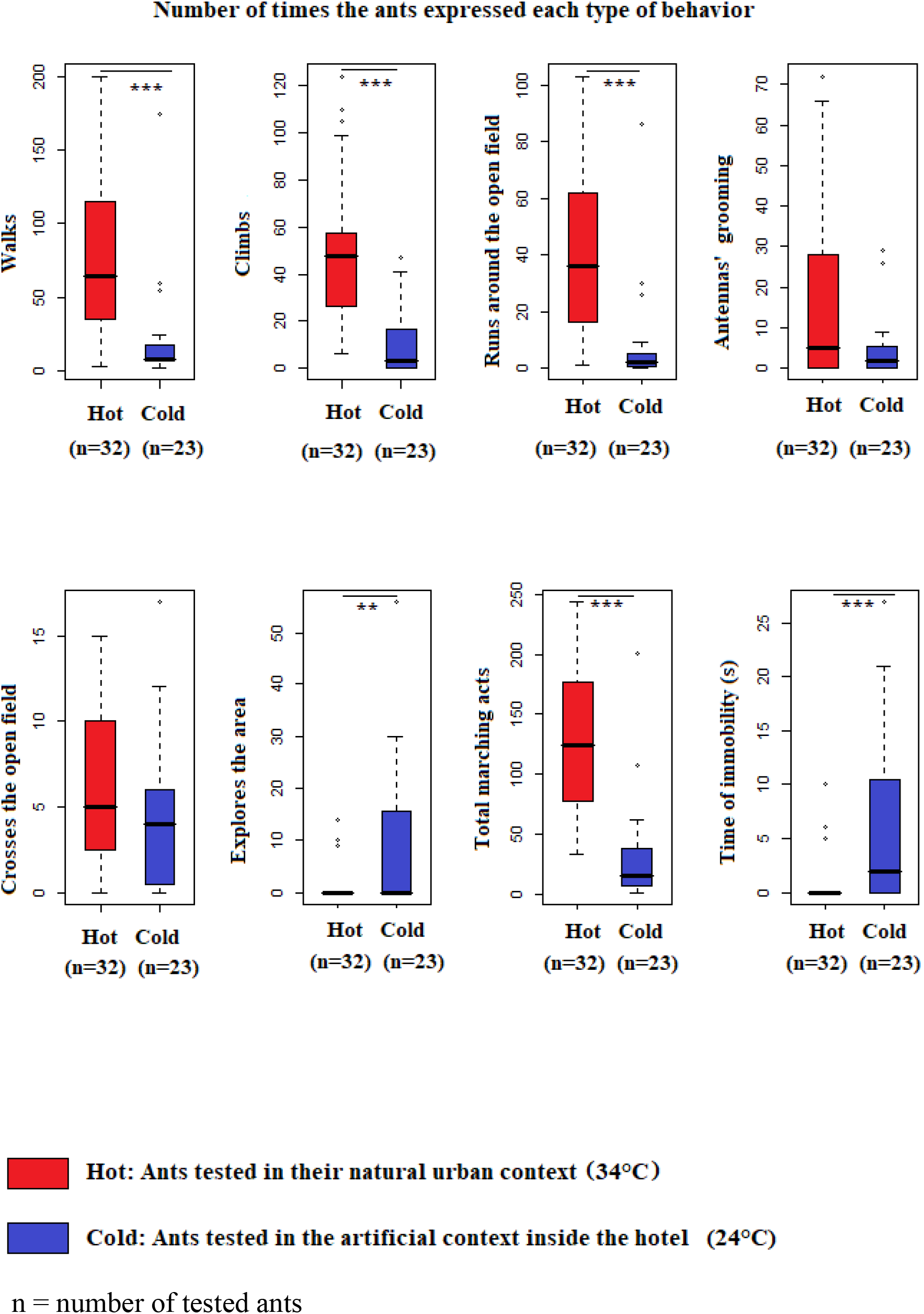
Comparison of *Camponotus atlantis* activity in the open field test when conducted in a hot environment and in a cold environment.

When I tested the ants at 34°C I did not observe any significant difference between the behaviour of individuals taken close from the nest and individuals taken far from the nest entrance (Mann Withney Test: p>0.05) **(Figure 10)**.

**Figure 10:**
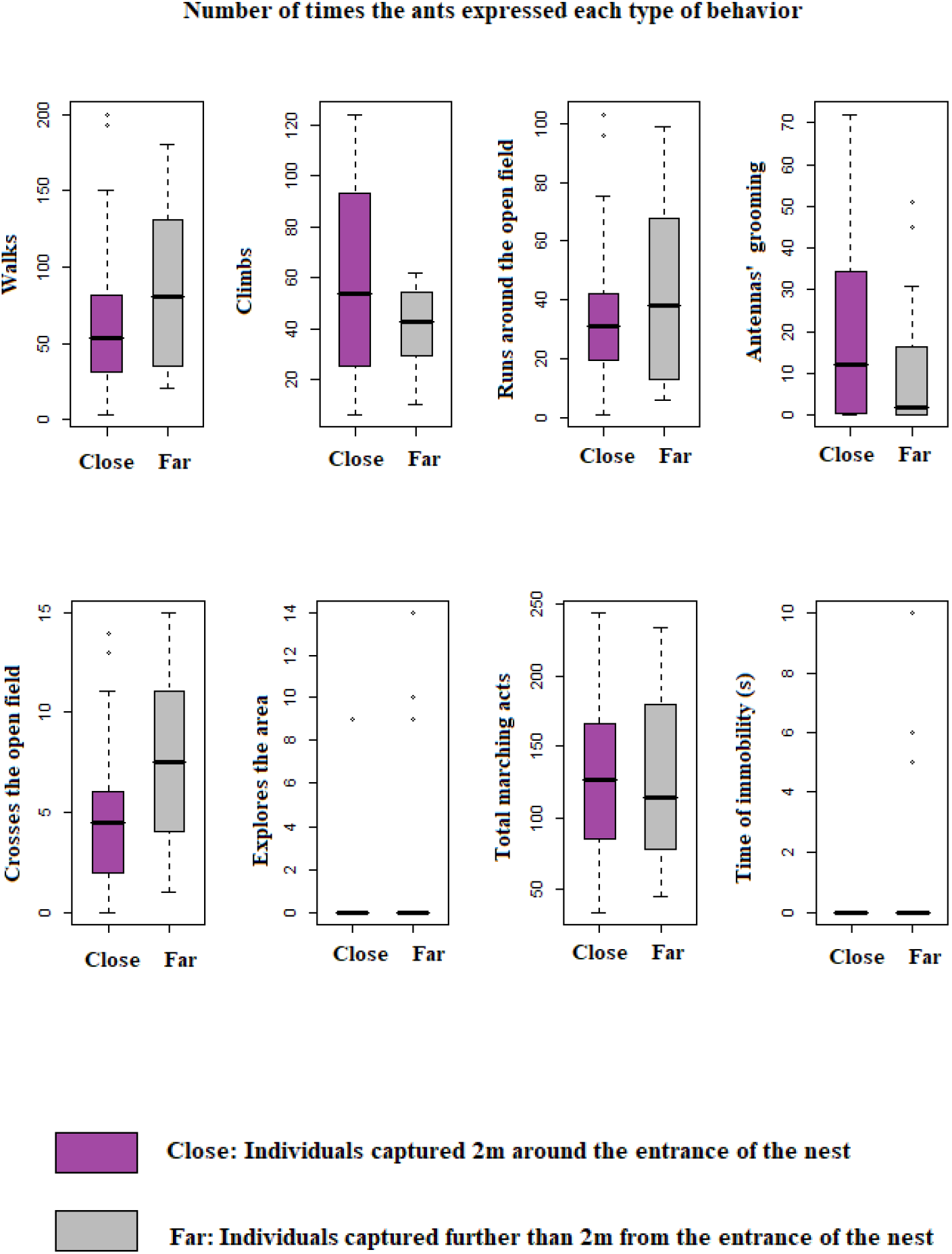
Comparison of *Camponotus atlantis* activity in a hot environment (34°C)

Whereas at 24°C individuals taken far from the entrance of the nest were significantly more active than individuals taken close from the nest entrance (they walked and run more frequently). On the contrary, individuals taken close from the entrance appeared to be more explorative than the others p<0.01 **(Figure 11)**.

**Figure 11:**
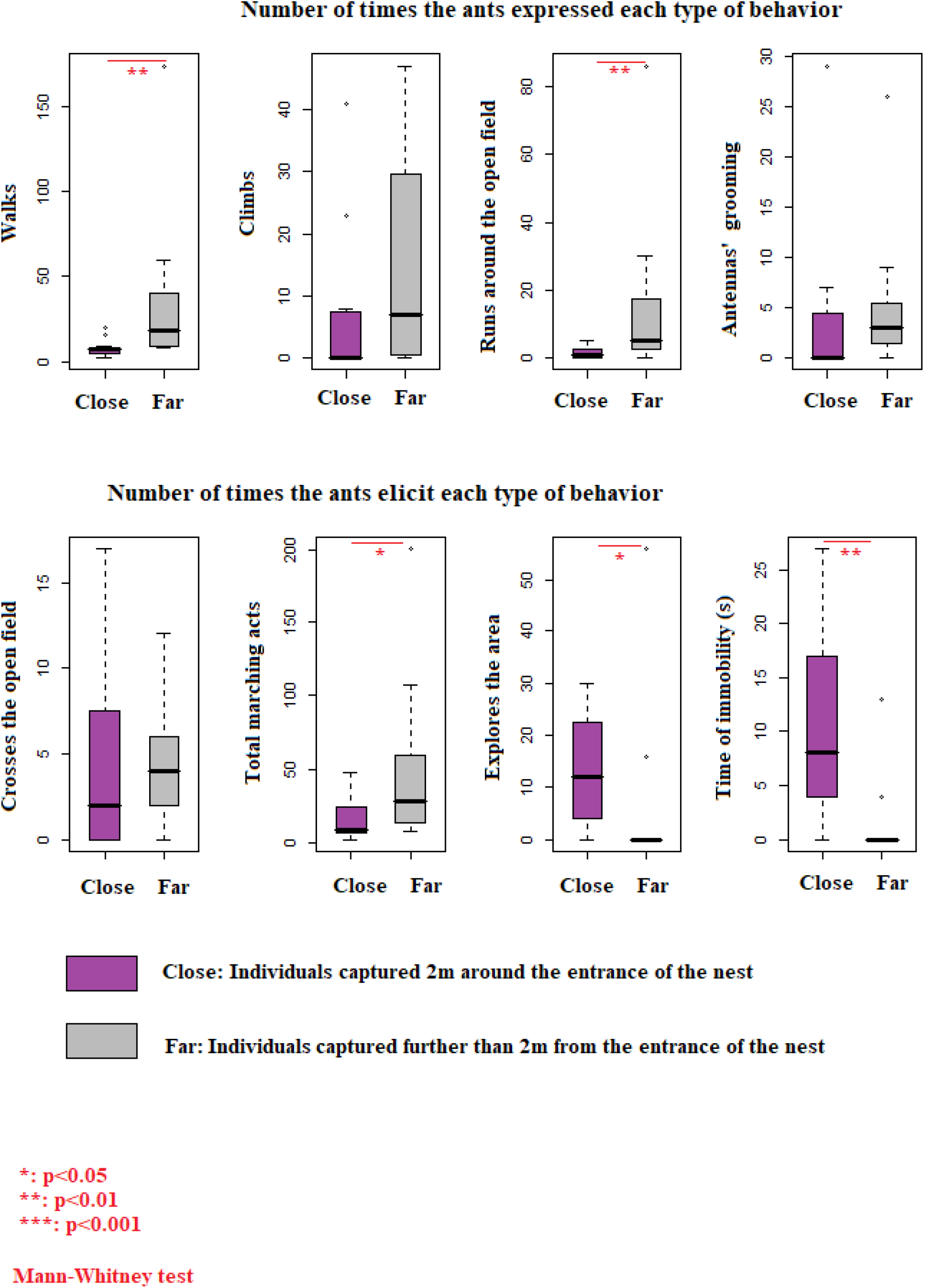
Comparison of *Camponotus atlantis* activity in a cold environment (24°C)

When I compared behavioural reactions between workers and soldiers in *Messor barbarus* group I observed that antenna grooming was a significantly more frequent behaviour in Soldiers than in Workers **(Figure 12)**.

**Figure 12:**
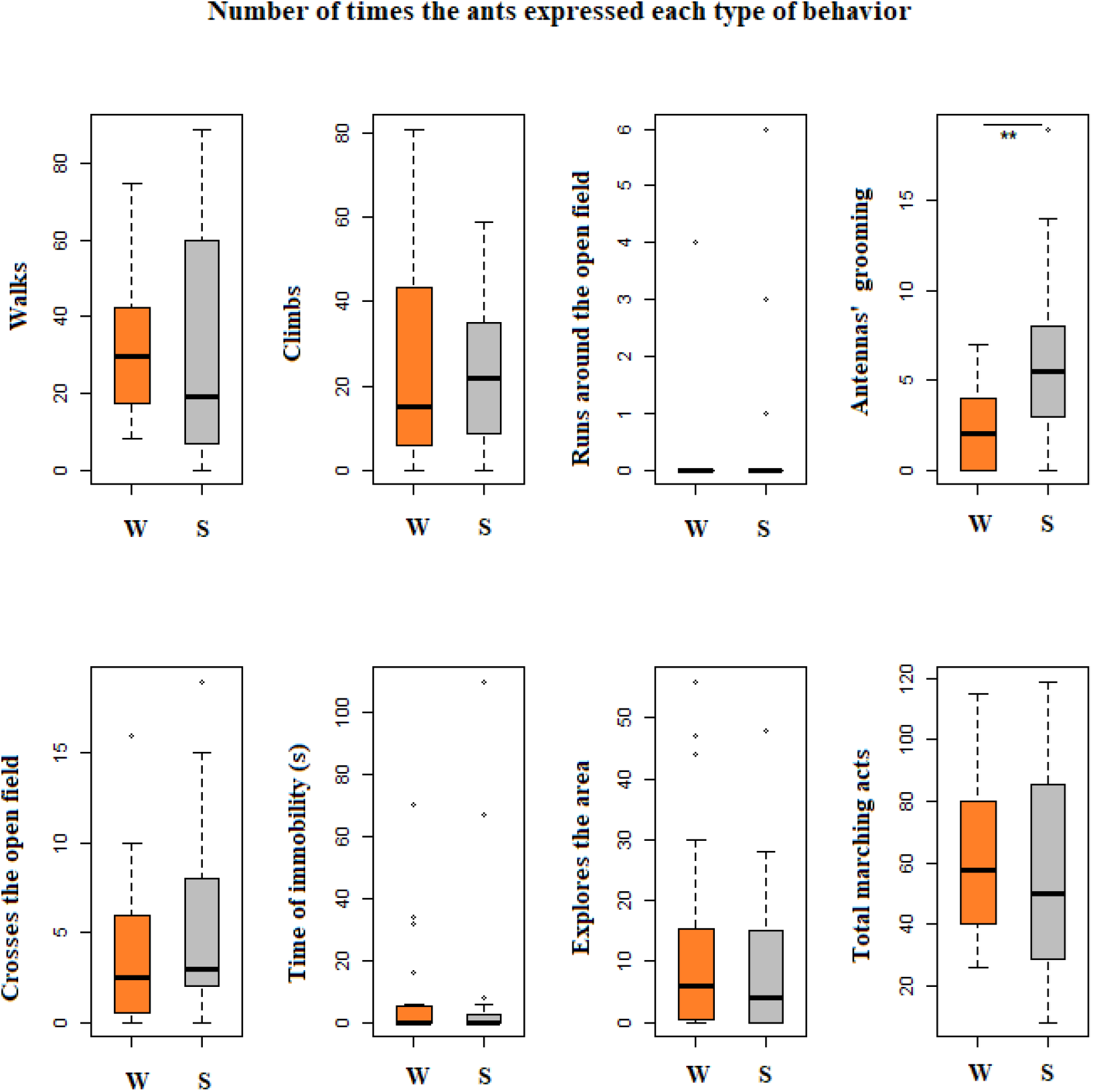
Comparison of behavioural reactions in *Messor barbarus* between Workers (W) and Soldiers (S) in the open field test.

In the group of *Atta cephalotes*, the soldiers crossed the open field and explored the area significantly more frequently than the workers **(Figure 13)**.

**Figure 13:**
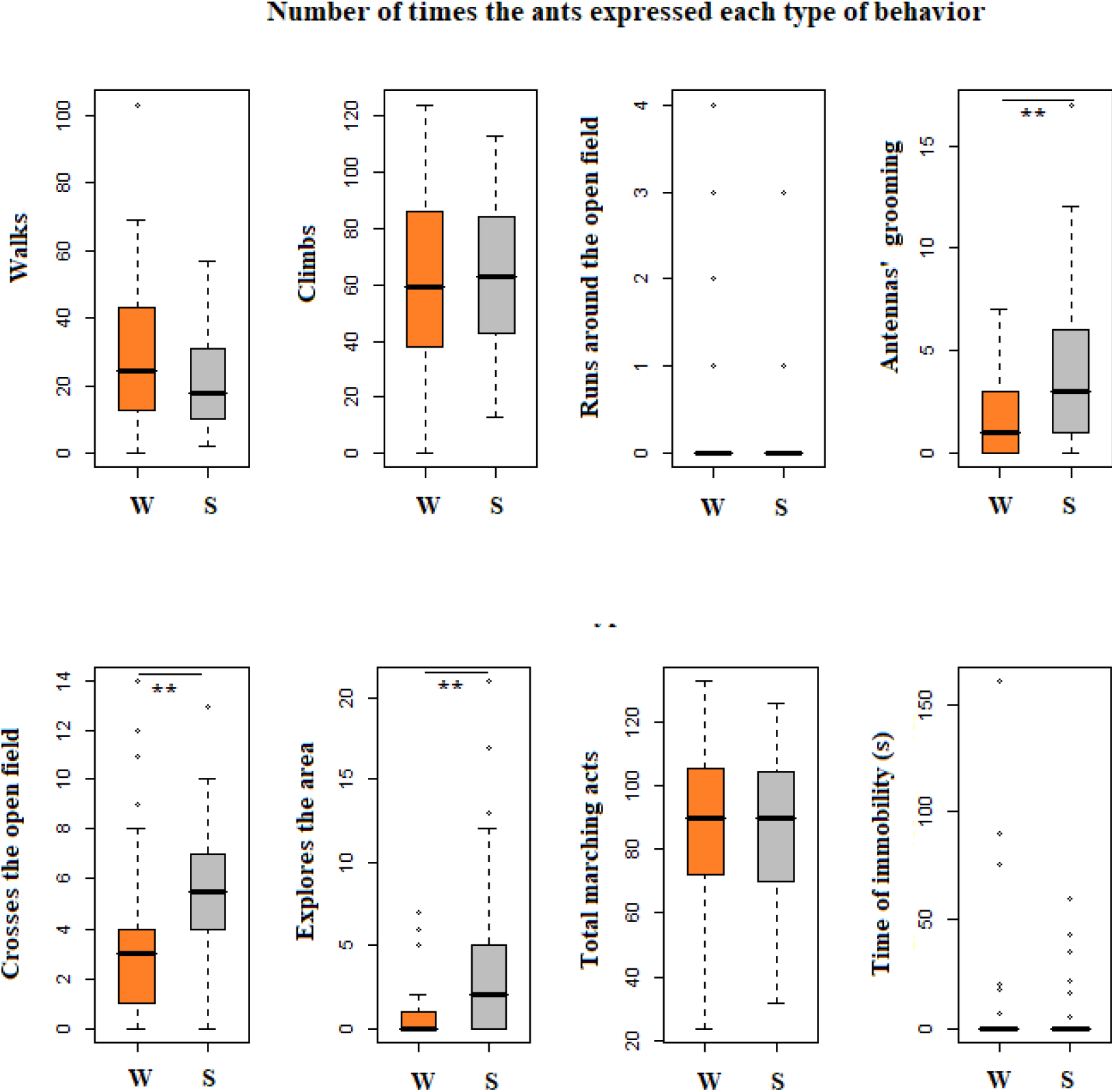
Comparison of behavioural reactions in *Atta cephalotes* between Workers (W) and Soldiers (S) in the open field test.

When comparing all the species in their response to open field test, we observed that invasive species that have colonized urban areas, islands and the rocky coast appeared to be bolder than the two species that harvest and cultivate the underground soil and get out from nests less frequently than the invasive ones. In fact, *Camponotus atlantis, Solenopsis invicta* and *Crematogaster crinosa* appeared to walk, climb the walls, cross the open field and run around the open field significantly more frequently than *Messor barbarus* and *Atta cephalotes*, indicating that the invasive species tend to be more hyperactive **(Figure 14)**.

**Figure 14:**
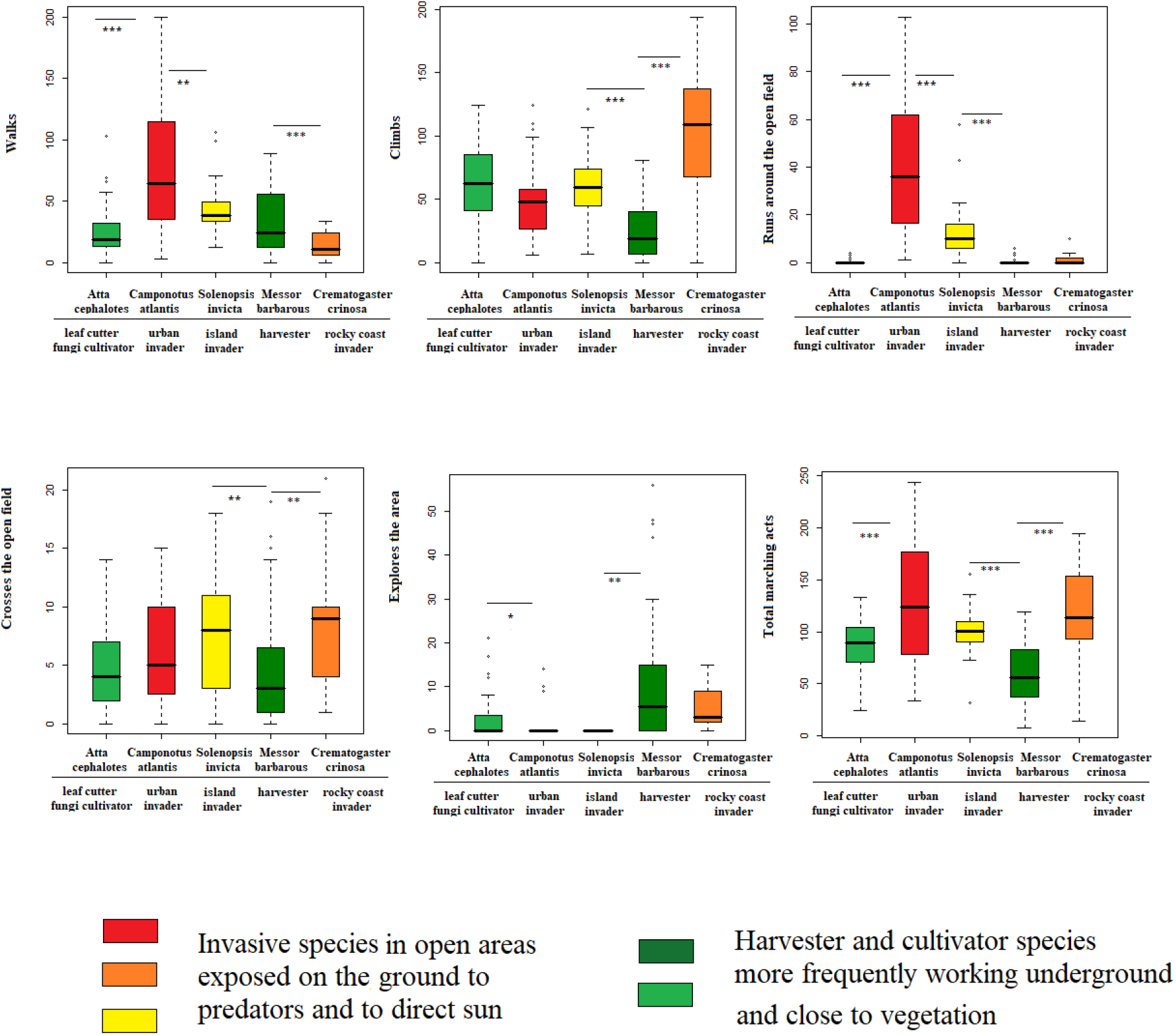
Comparison of open field test activity between the five studied species and their ecological contexts.

## Discussion

In this study I found for the first time that differences in neophobia in an open field test can be observed between ants from different species and between ants from different castes of a same species suggesting the existence of a behavioural diversity of responses to explorative tests.

I observed differences in individuals’ response to the open field test between ants that were captured close to the nest entrance and far from the nest entrance with a systematic result that indicates that individuals that forage far from the nest are bolder in open field test than individuals that forage close to the nest. My observations reveal the existence of two types of profiles (far explorers and close explorers) which is in agreement with other studies conducted on species studied on the field as it has been shown for example that in some ant species the old foragers explore further from the nest than younger ones or than nurses who leave less frequently the nest (Rodriguez et al. 2022, Gallotti and Chialvo 2023).

A relation between neophobia and dispersal has been observed in vertebrates like birds (Great tits, *Parus major*) and in terrestrial tortoises like Hermann tortoise, *Testudo hermanni*. Individuals that are bold in neophobia tests travel higher distances than shy ones in the wild (Digemanse et al.2003, Rodriguez et al. 2019). Moreover, Dingemanse et al. (2003) observed that fast exploring great tits had offspring that dispersed further. They also obtained that immigrant individuals arriving in a new habitat, were faster explorers in novel environment tests than locally born individuals. Here I observed for the first time that ant species may present individuals with different tendencies to disperse and that the ones that disperse the more are also bolder in open field test. The results of the comparison between species also suggest that they differ in their reactions towards a novel environment like the open field and that the species that tend to colonize more ecosystems and that are settled in open spaces like urban areas, islands and rocky landscapes that are more exposed to sun and to predators appeared to be bolder and more active in the open field test. We can think that there are behavioural syndromes like the ones describe by Sih and collaborators (2004) for crayfish or by Labaude et al. 2018 in beetles in another group of invertebrates, here the ants.

Jones and Gray observed behavioural differences in neophobia and open field test between males and females. In one hand, Jones (1977, 1982) observed that female chicks fed significantly sooner, longer and more than males when confronted to novel blue food and that females presented less behavioural inhibition when placed in a novel environment or in open field. In the same way, female rodents seem to explore sooner a novel environment than males (Gray 1971, 1974). In the other species like ungulates and primates, females appear to be more fearful than males (Buirski et al. 1978, Crepeau and Newman 1991). Here I compared two ant castes (soldiers and workers) in two different species of ants (*Messor barbarus* and *Atta cephalotes*) and I found for the two species that soldiers were bolder in the open field test than workers, they are more active (grooming and locomotor behaviours) and explore the open field more frequently than workers. Soldiers take more frequently the risk to cross the open field by leaving the “secure comfort” of glass walls than workers which seems to be in accordance with their social function in colonies.

Another result from this study is that activity in the open field tests increases with the ambient temperature during the test probably indicating that hot conditions elicit an increase in metabolic rate in ants provoking the display of more locomotor and exploration behaviours. It is known that invertebrates and in particular insects increase their activity with temperature (Abdullah 1961, Neven 2000).

The results obtained in this study suggest that ant species present personality different traits concerning boldness to explore novel environments. The fact that they can modulate their behaviour depending on temperature, the fact that individuals from a same species like far and close foragers or soldiers and worker can express different levels of fear reaction in the open field test suggest that there is a huge plasticity in explorative behaviour in this invertebrate taxon that may explain its success in so many habitats.

This hypothesis is supported by the observations concerning the different ecological realities of the studied species. Actually, we obtained that species living in exposed habitats like cities, rocky areas and islands are bolder and take more often risks to cross unknown environments than species that live in a very undergrounded environment and spend more time under the soil close to the protection of nests walls.

Different debates have been raised and major questions are still open concerning three essential points. One first question is if there is constancy or repeatability of interindividual differences across the time (Jones 1977). In one hand, constancy across the time of interindividual differences is generally interpreted as the existence of temperaments or personalities (Gosling 2001). On the other hand, the absence of consistency is interpreted as the expression of context or state dependent behaviours (Van Oers et al. 2005) or as the expression of behavioural plasticity (as the individual can behave in one way one time and another way another time) (Pfennig et al. 1993, Neff and Sherman 2003). More studies on invertebrate interindividual differences must be conducted in order to know if they are really different personality stable traits.

Greenberg (2003) proposed that the primary benefit of neophobia is protection against danger (the Dangerous Niche Hypothesis). Neophobia can be driven by natural selection based on environmental patterns because neophobia can have a strong innate (genetic) component (Mettke-Hofmann, 2017). However, individual neophobic responses are also shaped by ontogenetic experience, which can result in more or less neophobia (phenotypically plastic neophobia: Brown et al. 2013). More studies should be conducted in order to better understand how this neophobia may be modulated in space and time during the evolution of each ant species and how it can explain the amazing adaptations of ants to several ecosystems.

## References

Abdullah M. 1961. Behavioural effects of temperature on insects. Ohio J. Sci., 61 (1961), p. 212

Bates, JE. 1986. The measurement of temperament. In Plomin, R. & Dunn, J. (Eds.), The study of temperament: Changes, continuities and challenges (pp. 1–11). Hillsdale, NJ: Lawrence Erlbaum.

Bates, J. E. (1989). Concepts and measures of temperament. In G. A. Kohnstamm, J. E. Bates, & M. K. Rothbart (Eds.), Temperament in childhood (pp. 3–26). Oxford, England: John Wiley & Sons.

Bolton, 1973. The ant genera of West Afruca: a synonymic synopsis with keys (Hymenoptera: Formicidae). Bulletin of the British Museun (Natural History). Entomology 27: 317–368.

Brown GE, Chivers DP. 2005. Learning as an adaptive response to predation. In Ecology of Predator - Prey Interactions (eds Barbosa and I Castellanos). Pp. 34–54. Oxford University Press.

Brown GE, Ferrari M, Elvidge C, Ramnarine I, Chivers D. 2013. Phenotypically plastic neophobie : a response to variable predation risk. Proceedings of the Royal Society B : Biological Sciences 280

Buirski P, Plutchik R, Kellerman H. 1978. Sex differences, dominance, and personality in the chimpanzee. Animal behaviour 26 (1): 123–129

Candler S, Bernal X. 2015. Differences in neophobia between cane toads from introduced and native populations. Behavioural Ecology 26: 97–104

Carazo P, Noble D, Chandrasoma D, Whiting MJ. 2014. Sex and boldness explain individual differences in spatial learning in a lizard. Proceedings of the Royal Society 281 (1782):

Carter J, Heinsohn R, Goldizen A, Biro P. 2012. Boldness, trappability and sampling bias in wild lizards. Animal Behaviour 83 (4): 1051–1058

Chapple D, Simmonds S, Wong B. 2012. Can behavioral personality traits influence the success of unintentional species introductions? Trends in Ecology and Evolution 27: 57–64

Chitkka A, Wurm Y, Chitkka L. 2012. Epigenetics: the making of ant castes. Current Biology 9:R835–R838 https://doi.org/10.1016/j.cub.2012.07.045

Coleman, K., Wilson D.S. 1998. Shyness and boldness in pumpkinseed sunfish: individual differences are context-specific. Animal Behaviour 56: 927–936

Corrêa M, Bieber A, Wirth R, Leal I. 2005. Occurrence of Atta cephalotes inAlagoas, northeastern Brazil. Neotropical Entomology 34: 695–698

Crepeau, L. J., & Newman, J. D. (1991). Gender differences in reactivity of adult squirrel monkeys to short-term environmental challenges. Neuroscience and Biobehavioral Reviews, 15(4), 469– 471. https://doi.org/10.1016/S0149-7634(05)80133-2

Detrain, C., et al. (2000). A field assessment of optimal foraging in ants: Trail patterns and seed retrieval by the European harvester ant Messor barbarus. Insectes Sociaux 47, 56–62.

Dingemanse N, Both C, Van Noordwijk A, Rutten A, Drent P. 2003. Natal dispersal and personalities in Great tits. Proceedings of the Royal Society B 270: 741–747

Fox R, Millam J. 2007. Novelty and individual differences influence neophobia in orange-winged Amazon parrots (Amazona amazonica). Applied Animal Behavior Science 104: 107–115

Galib S, Sun J, Twiss S, Lucas M. 2022. Personality, density and habitat drive the dispersal of invasive crayfish. https://doi.org/10.1038/s41598-021-04228-1

Gallotti R, Chialvo D. 2018. How ants move: collective versus individual scaling properties. Journal of the Royal Society Interface 15 : DOI:10.1098/rsif.2018.0223

Hita Garcia F, Wiesel E, Fisher G. 2013. The ants of Kenya (Hymenoptera : Formicidae) faunal overview, first species checklist, bibliography, accounts for all genera, and discussion on taxonomy and zoogeography. Journal of East Anfrican Natural Histoty 101 (2) : 127–122

Goldsmith, H.H, Buss, AH, Plomin R, Rothbart MK, Thomas A, Chess S, Hinde S, McCall RB. 1987. What is temperament? Four approaches. Child Development 58 (2): 505–529

GoslingSD. 2001. From mice to men: what can we learn about personality from animal research? Psychological Bulletin 127 (1): 45–86

Gray JA. 1971. Sex differences in emotional behaviour in mammals including man: endocrine bases. Acta Psychologica 35(1):29–46

Gray JA, Lalljee B. 1974. Sex differences in emotional behavior in the rat. Correlation between open field defecation and active avoidance. Animal Behaviour 22: 856–861

Greenberg R. (2003). The role of neophobia and neophilia in the development of innovative behaviour of birds. In Animal Innovation (eds. SM Reades and KN Laland) pp. 175–196 Oxford University Press. New York

Helmes, IV, J.A., Bridge, E.S. 2017. Range expansion drives the evolution of alternate reproductive strategies in invasive fire ants. NeoBiota, 33: 67–82.

Jones B.R. 1977a. Sex and strain differences in the open-field responses of the domestic chick. Applied Animal Ethology 3: 255–261

Jones B.R. 1982. Effects of early environmental enrichment upon open-field behaviour and timidity in the domestic chick. Developmental Psychobiology 15: 105–111

Labaude S, Griffin C, O’Donnell N. 2018. Description of a personality syndrome in a common and invasive ground beetle (Coleoptera: Carabidae). Scientific Reports Nature Portfolio 8 : DOI:10.1038/s41598-018-35569-z

Lee D.N., Tang-Martinez Z. 2009. What behavioral syndrome ? Individual differences and the search for exploratory behavioural profiles among prairie voles. XXXI° International Ethological Conference 19-24 August 2009. Rennes France

Le Scolan, N., Hausberger, H., Wolff, A. 1997. Stability over situations in temperamental traits of horses as revealed by experimental and scoring approaches. Behavioural Processes 41: 257–266

Mettke-Hofmann C, Ebert C, Schmidt T, Steiger S, Stieb S. 2005. Personality Traits in Resident and Migratory Warbler Species. Behaviour 142 (9): 1357–1375

Mettke-Hofmann, C (2017). Neophobia. In Encyclopedia of Animal Cognition and Behavior (eds J. Vonk and T Shackelford), pp 1-8. Springer Internashional Publishing)

Montoya-Lerma J, Ulloa Chacón P, Manzano M. 2006. Characterization of the nest of the leaf-cutting ant Atta cephalotes in Cali (Colombia). Entomología 32: 151–158

Neff, B.D., Sherman, P.W. 2004. Behavioral syndromes versus darwinian algorithms. Trends in Ecology and Evolution 19: 621–622

Neven, L.G., 2000. Physiological responses of insects to heat. Postharvest Biol Technol 21, 103 – 111. doi:10.1016/S0925-5214(00)00169-1

Pfennig, D.W., Reeve, H.K., Sherman P.W. 1993. Kin recognition and cannibalism in spade foot toad tadpoles. Animal Behaviour 46: 87–94

Rodríguez A, Caron S, Ballouard JM. Differences in personality / temperament in the endangered tortoise Testudo hermanni and their implications on dispersal and conservation https://doi.org/10.1101/2021.08.05.45529

Characterization of an ant colony (Paraponera clavata) during rest and relocation phases: an experimentally induced protocol

Sih A, Kats LB, Maurer EF. 2003. Behavioural correlations across situations and the evolution of antipredator behaviour in a sunfish–salamander system. Animal Behavior 65: 29–44

Sih A, Bell A, Johnson JC. 2004. Behavioral syndromes: an ecological and evolutionary overview. Trends in Ecology and Evolution 19: 372–378

Sinn D, Gosling S, Moltschaniwskyj N. 2008. Development of shy/bold behaviour in squid: context-specific phenotypes associated with developmental plasticity. Animal Behaviour 75: 433–442

Taiwo A. 2015. The open field and animal behaviour. Bc Tesis.

Tremmell M, Müller C. 2013. Insect personality depends on environmental conditions. Behavioral Ecology 24 (2) : 386–392

Van Oers, K., Klunder, M., Drent P. 2005. Context dependence of personalities: risk-taking behavior in a social and nonsocial situation. Behavioural Ecology 16: 716–723

Westerman PR, Atanackovic V, Royo-Esnal A. Torra J. 2012. Differential weed seed removal in dryaland cereals. Arthropod-Plant Interaction 6:591–599

